# Diverse arsenic-containing lipids in the surface ocean

**DOI:** 10.1101/2021.03.22.436501

**Authors:** Katherine R. Heal, Ashley E. Maloney, Anitra E. Ingalls, Randelle M. Bundy

## Abstract

Arsenic is present at nanomolar levels throughout the surface ocean, and microbes assimilate this toxic element due to its similarity to inorganic phosphorus. Although the concentration and characterization of dissolved arsenic has been a focus of ocean studies, the size of the particulate arsenic pool and its partitioning into organic molecules within the microbial community is not known. We measured the particulate pool of arsenic in five surface samples from the open ocean and determined the contribution of arsenic-containing lipids to this pool. Here we show that the accumulation of arsenic into lipids is a widespread phenomenon in the surface ocean. Particulate arsenic concentrations were 15 to 42 pmol L^−1^ with 7–20% of the particulate arsenic pool in the form of arsenolipids. We characterized these arsenolipids and found that arsenosugar phospholipids dominated the arsenolipid pools in our samples with a minor component of arsenohydrocarbons and other unidentified lipids. A significant portion of the arsenosugar phospholipids (up to 35%) were present as previously undescribed mixed acyl ether lipids, suggesting a bacterial source.

**Scientific significance statement:** Marine microbes experience constant exposure to the toxic element arsenic. However, there are no baseline measurements of how much arsenic accumulates in microbial communities nor do we know the full spectrum of arsenic containing biomolecules produced in marine systems. In culture-based studies there is strong evidence that phytoplankton synthesize arsenic-containing lipids, but these lipids have not been observed in natural communities of marine microbes. Here we make measurements of bulk particulate arsenic at five sites in the surface open ocean and show that a significant portion of this particulate arsenic is present as complex arsenic-containing lipids. We characterize this arsenolipid pool chemically and quantitatively to show a variety of chemically distinct and quantitatively significant lipids that expand our understanding of marine arsenic biogeochemistry.

**Data availability statement:** Mass spectrometry data and its associated metadata will be available on Dryad with publication. Scripts for data processing and figure generation found on github at https://github.com/kheal/particulate_As_data_analysis.

## Introduction

Arsenic is present at nanomolar levels throughout the world’s oceans. Its nutrient-like profile indicates accumulation within surface ocean marine microbial communities and subsequent export via sinking particles into the deeper ocean (Middelburg et al. 1988; Cutter and Cutter 1995). Oceanographic studies of arsenic have focused primarily on the quantification and characterization of the dissolved species of arsenic in seawater. These studies have shown that within the dissolved pool, simple organic arsenic compounds like mono- and dimethylarsenic acid build up in nutrient depleted areas of the ocean, particularly regions with low phosphorus (Andreae 1978; Cutter et al. 2001; Wurl et al. 2013). The distribution of reduced and methylated arsenic species in seawater suggests that arsenic biotransformations occur primarily in low phosphorus environments to detoxify arsenate (AsO_4_^3−^) which microbes accidentally take up due to its similarity to phosphate (PO_4_^3−^).

Although oceanographic studies of arsenic often focus on the element’s toxicity, arsenic can have a much richer set of roles within microbial systems. For instance, some bacteria can use AsO_4_^3−^ as an electron acceptor for anaerobic respiration or arsenite (AsO_3_^3−^) as an energy yielding oxidant (Mazumder et al. 2020), and these processes have been inferred to exist in modern marine systems from gene expression studies (Saunders et al. 2019). In picocyanobacteria, the genetic potential for arsenic methylation (arsenic methyl transferase, *asrM*) is more common than the genetic potential for the ‘reduce and efflux’ arsenic detoxification mechanism (Saunders and Rocap 2016), and *asrM* is widespread among other marine phytoplankton clades (Chen et al. 2017). In coastal ecosystems, macro and microalgae transform arsenic into organic-arsenic compounds well beyond what one would expect as a detoxification mechanism (Vinogradov 1953; Lund 1973; Sanders 1979). These organic arsenic compounds accumulate in fish (i.e. Taleshi et al. 2014) and are produced by monocultures of green algae, diatoms, and cyanobacteria in laboratory settings (Duncan et al. 2013; Xue et al. 2014; Řezanka et al. 2019). Arsenolipids have also been observed in lake sediments and lacustrine suspended particles (Glabonjat et al. 2019, 2020). Together, these findings suggest that the production of organic arsenic compounds is a widespread process in aquatic settings. In this work, we quantify the stock of arsenic associated with microbial biomass and evaluate whether natural mixed microbial communities in the surface ocean harbor arsenolipids.

## Methods

### Overview

Surface samples for particulate arsenic speciation and quantification were collected from five locations noted in Table 1 (more details in Table S1). These samples include two from oligotrophic subtropical regions: one from the North Pacific (ALOHA: A Long-Term Oligotrophic Habitat Assessment) and one from the North Atlantic (BATS: Bermuda Atlantic Time Series). Three additional samples were collected from the equatorial upwelling influenced Eastern Tropical North Pacific (ETNP), including two offshore samples (ETNP-PS1 and ETNP-PS2) and one coastal sample (ETNP-PS3). We took quantitative subsamples for total particulate carbon and total particulate arsenic. We extracted samples for arsenolipids which we analyzed by liquid chromatography – inductively coupled plasma mass spectrometry (LC-ICP-MS, for quantification) and liquid chromatography – high resolution electrospray ionization mass spectrometry (LC-HR-ESI-MS, for identification of individual lipids). Details for each step are in Supplemental Methods.

**Table 1:** Summary of quantitative results of particulate arsenic and arsenolipids. Standard deviations are presented for bulk particulate As and C measurements. Bulk particulate arsenic measurements are mean and standard deviation from a single filter; particulate C measurements are mean and standard deviation from multiple filters.

| Sample | Latitude | Longitude | particulate As (pM) | particulate C (μM) | particulate As/C (x10<sup>6</sup>) | total As lipids (pM) | particulate As as lipids (%) | AsHC (pM) | AsSugPL (pM) | AsSugPeL (pM) |
| --- | --- | --- | --- | --- | --- | --- | --- | --- | --- | --- |
| ALOHA | 22.75 | -158 | 36 ± 8.7<br>(n = 3) | 2 ± 0.3<br>(n = 14) | 18 ± 5.1 | 4.9 | 10 | 0.064 | 0.76 |  |
| BATS | 31.66 | -64.167 | 42 ± 10<br>(n = 3) | 3.3 ± 1<br>(n = 3) | 13 ± 5 | 2.9 | 7 | 0.083 | 0.38 |  |
| ETNP-PS1 | 10 | -113 | 15 ± 0.75<br>(n = 3) | 3.7 ± 1.1<br>(n = 6) | 4.1 ± 1.2 | 2.6 | 20 | 0.12 | 0.47 |  |
| ETNP-PS2 | 15.77 | -103 | 19 ± 1.3<br>(n = 3) | 3 ± 0.4<br>(n = 3) | 6.3 ± 0.94 | 3.5 | 20 | 0.14 | 0.55 | 0.19 |
| ETNP-PS3 | 17.68 | -102.35 | 33 ± 3.9<br>(n = 3) | 9.5 ± 5.1<br>(n = 7) | 3.4 ± 1.9 | 5.7 | 20 | 0.16 | 0.74 | 0.4 |

### Total particulate carbon

Subsamples of known surface area were taken from filters and dried overnight at 60 °C. Dried samples were pelletized in 9 x10 mm tin capsules (Elemental Microanalysis and Costech) and run on a vario ISOTOPE select (Elementar) CHNOS Elemental Analyzer to quantify total C. Measurements were calibrated with an in-house aminocaproic acid standard (ACROS).

### Total particulate arsenic

Three quantitative subsamples from each sample were taken and digested for four hours in 1 mL of 50% HNO_3_ at 110 °C (Planquette and Sherrell 2012). We reconstituted the dried digests in 1 mL of 2% HNO_3_, 2.5% MeOH, 1 ppb Rh in MQ water. We analyzed concentrations of these extracts on a Thermo quadrupole ICP-MS (iCAP-RQ, Thermo Scientific) paired with a Thermo Dionex 3000 HPLC used as an autosampler (no column), monitoring for ^75^As, ^77^Se (to monitor for ^35^Cl^40^Ar interference), ^78^Se, and ^103^Rh. We performed sample analyses in kinetic energy discrimination (KED) mode with He collision gas at 2.0 L min^−1^ which eliminated ^35^Cl^40^Ar interference.

### Arsenolipid analyses

We separated water-soluble compounds from lipid-soluble compounds to yield a crude lipid extract which we analyzed for total extractable arsenolipids. We performed a silica-gel clean up step as previously described (Glabonjat et al. 2014) that selects for arsenolipids. We used the same LC configuration for both LC-ICP-MS (for quantification) and LC-HR-ESI-MS (for identification) so peaks could be aligned. We tested multiple gradients with our ALOHA sample to ensure that the full ^75^As signal eluted within our chromatographic parameters and ran blanks between each sample. In brief, our LC set up was a Dionex Ultimate 3000 bio-inert LC equipped with a C18 column (ZORBAX SB-C18 by Agilent, 0.5×150 mm, 5 *µ*m particle size, 80 Å pore size) with a flow rate of 40 *µ*L min^−1^ and a gradient of a mixture of water and isopropyl alcohol, with 1% formic acid.

For quantification of arsenolipids, we injected crude lipid extracts and directed the flow from LC to the ICP-MS and monitored the ^75^As trace over time. As there are no commercial standards for arsenolipids, to quantify we used a proxy standard that was retained on our C18 column (cyanocobalamin), using an approach similar to what has been used to quantify nickel-bound ligands in seawater (Boiteau et al. 2016). We quantified each chromatographically resolved peak from the baseline (as calculated from the baseline (Liland et al. 2010)) and also quantified the total ^75^As signal retained on the LC column (not including the ^75^As signal associated with unretained compounds in the first five minutes).

For identification of arsenolipids, we injected both the crude and silica-gel cleaned extracts, directing the flow of the LC to the hybrid quadrupole Orbitrap (Q-Exactive HF, Thermo Scientific) for HR-ESI-MS. We collected both MS^1^ and MS^2^ data using data-dependent acquisition. We searched the resulting MS^1^ and MS^2^ scans for masses (MS^1^) or fragments (MS^2^) associated with arsenolipids from an in-silico database of arsenolipids curated in house from general structures previously reported (summarized in Table S2, database supplied in Tables S3 and S4).

## Results

### Bulk quantification of total arsenic and arsenolipids in marine suspended material

Total particulate arsenic ranged from 15–42 pM and the highest arsenic and As:C ratios were in the oligotrophic samples (HOT and BATS, Table 1). Peaks in ^75^As in the LC-ICP-MS chromatogram suggested the presence of individual arsenolipids that could be probed for identification (Figure 1). By integrating the total signal in the LC-ICP-MS traces (excepting small peaks near the start of the run associated with compounds not retained on the LC column), we estimated the concentration of total extracted arsenolipids to be 2.6–5.7 pM, or 7–20% of the total particulate arsenic pool in our five samples (Table 1).

**Figure 1:**
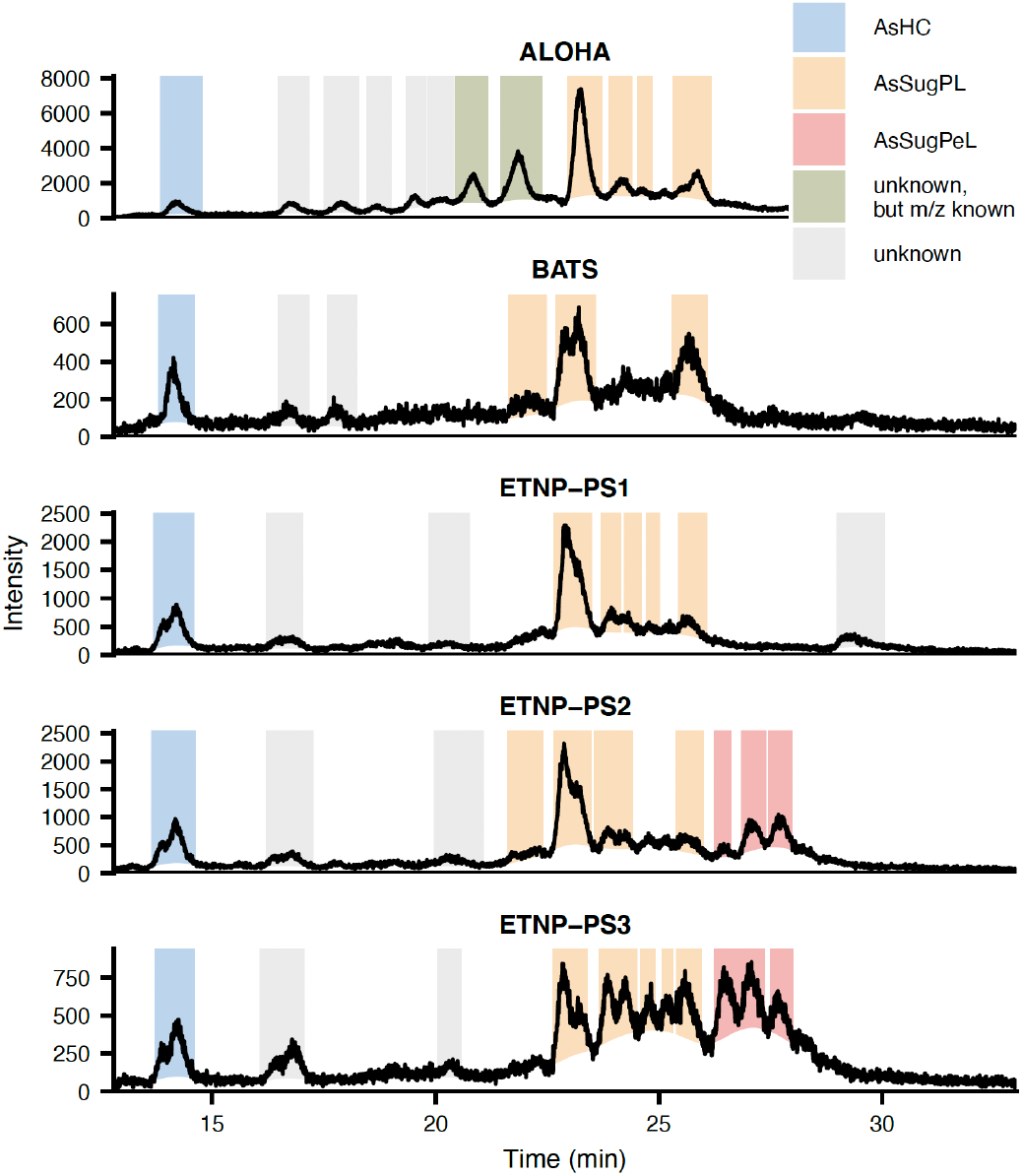
LC-ICPMS traces of the ^75^As intensity over the course of the chromatographic run in each of the five samples. Colored regions are individual lipids that were quantified, with identifications noted by different shades. There are no significant peaks before the time shown except some ^75^As signal associated with water soluble species that elutes with the solvent front. Note that different amounts of material extracted accounts for much of the difference in intensities, details in Table S1.

### Identification of organic-extractable arsenic compounds

Distinct peaks in the LC-ICP-MS ^75^As chromatograms in each of our samples corresponded to individual arsenolipids when aligned with the LC-HR-ESI-MS data. All samples showed a peak early in the chromatogram at approximately 14 minutes (Figure 1), which we identified as an arsenohydrocarbon (AsHC) with an *m/z* of 333.2139 (AsHC332, Figure 2), confirmation based on the MS^2^ fragmentation spectra (Figure 3A). In all samples, we saw peaks at 23 and 26 minutes in the LC-ICP-MS chromatograms which we identified as diacyl glycerol arsenosugar phospholipids (AsSugPLs) with *m/z*’s of 955.4893 (AsSugPL954) and 1011.5519 (AsSugPL1010). These masses were also associated with MS^2^ spectra showing diagnostic fragments for AsSugPLs including a 409.0245 fragment corresponding to the phosphorylated arsenosugar with a glycerol headgroup (C_10_H_23_AsO_10_P, Figures 2, 3B, 3C), as seen previously (i.e. Raab et al. 2013). More AsSugPLs including 902, 928, 930, 952, 956, 958, 982, and 984 were seen between 22 and 25 minutes in some of the samples (confirmed by MS^1^ and MS^2^ including the diagnostic 409.024 fragment) sometimes corresponding to smaller quantifiable ^75^As LC-ICP-MS peaks (Figure 4).

**Figure 2:**
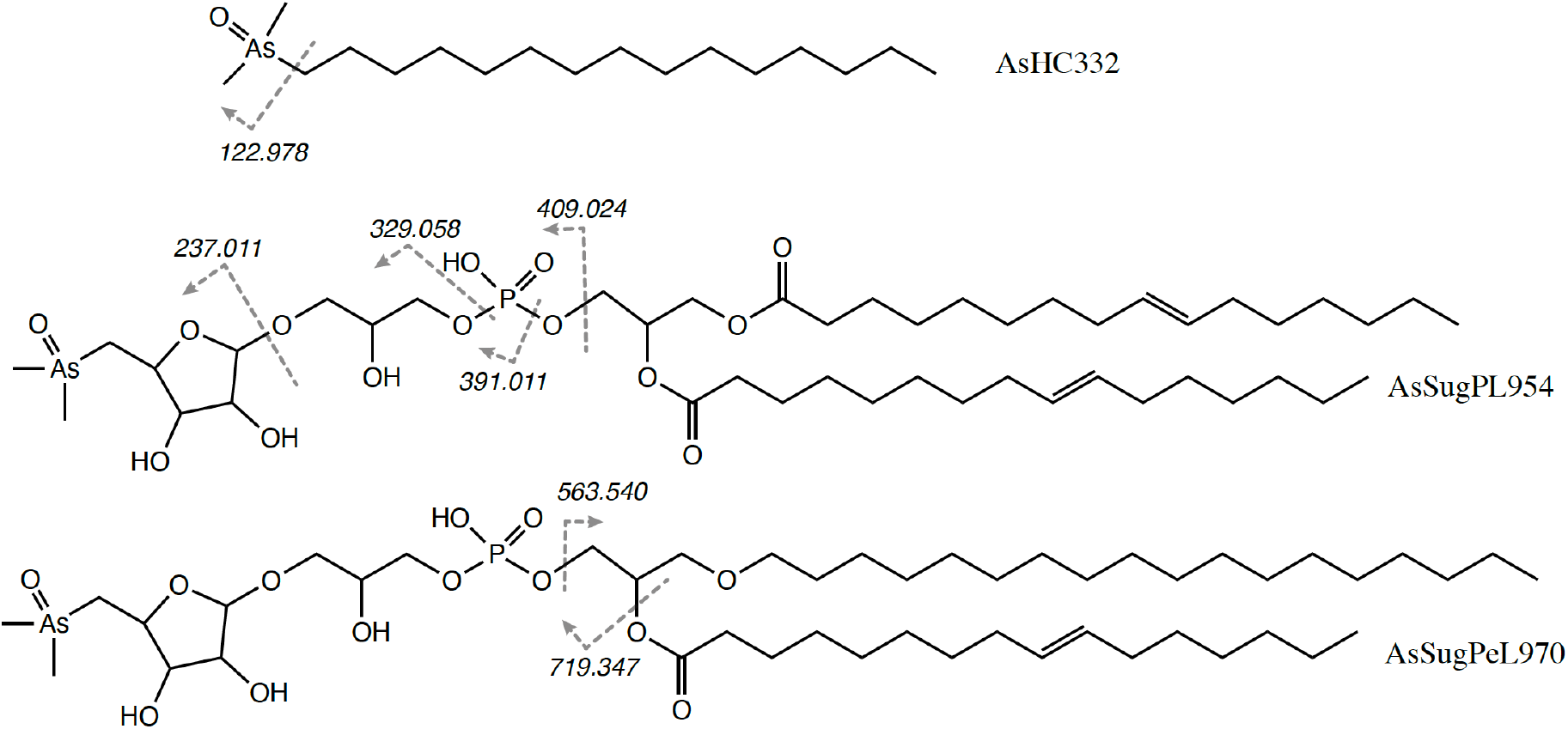
Arsenolipid structures. Note that just one example of the possible chains are shown. For AsSugPL we are not able to resolve individual chain length or saturation location, for AsSugPeL, chain lengths shown are derived from MS^2^ fragmentation, but saturation location on fatty acid chain is uncertain.

**Figure 3:**
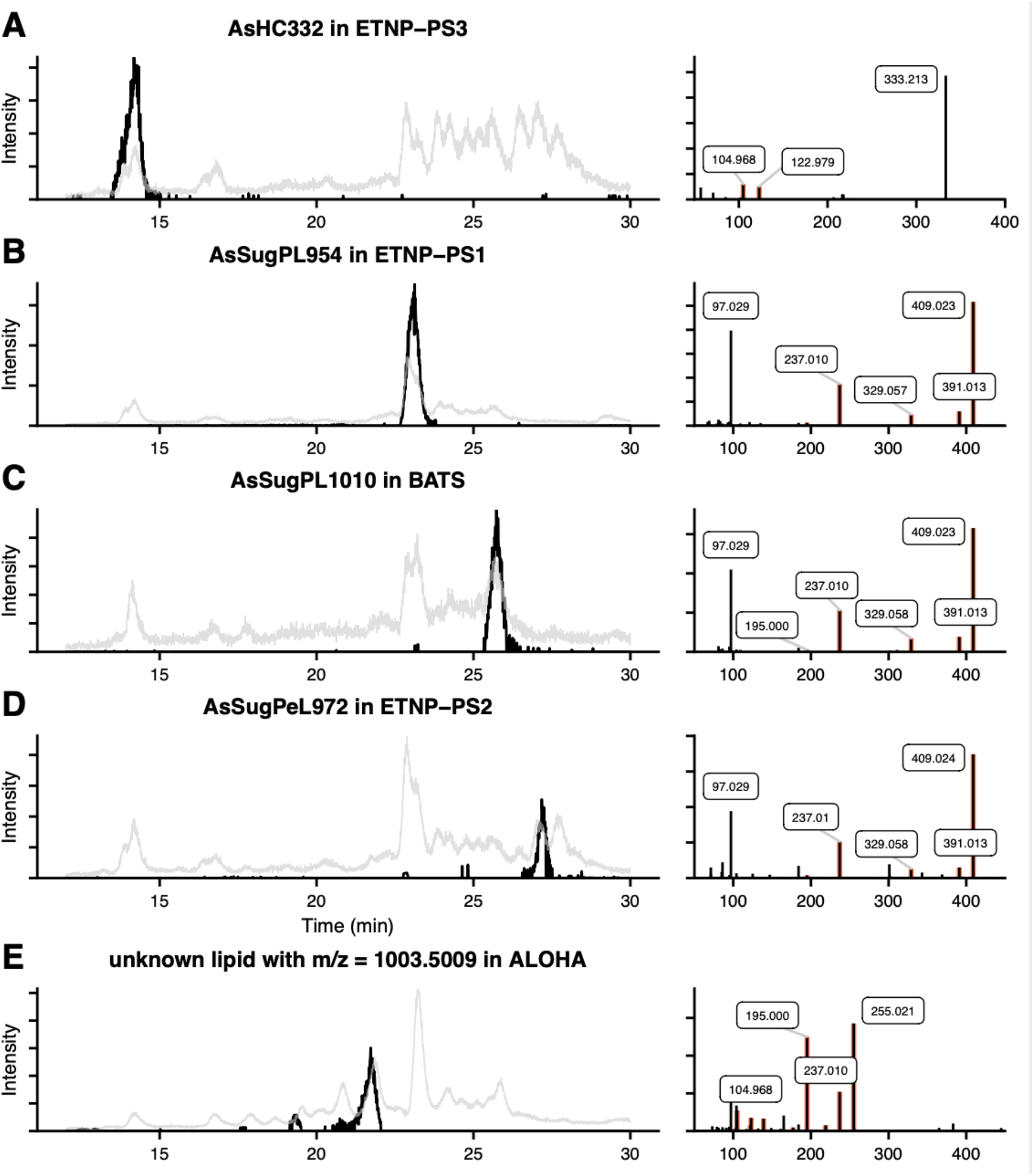
Identification of individual arsenolipids. Left panels show extracted ion chromatograms for identified lipids or masses from the LC-HR-ESI-MS (in black) overlaid on the LC-ICP-MS ^75^As signal (in grey). Right panels show MS^2^ spectra, the masses with arsenic highlighted – full observed MS^2^ for unknown lipids in ALOHA samples and AsSugPeLs, shown in Figures S1 and S2 respectively.

**Figure 4:**
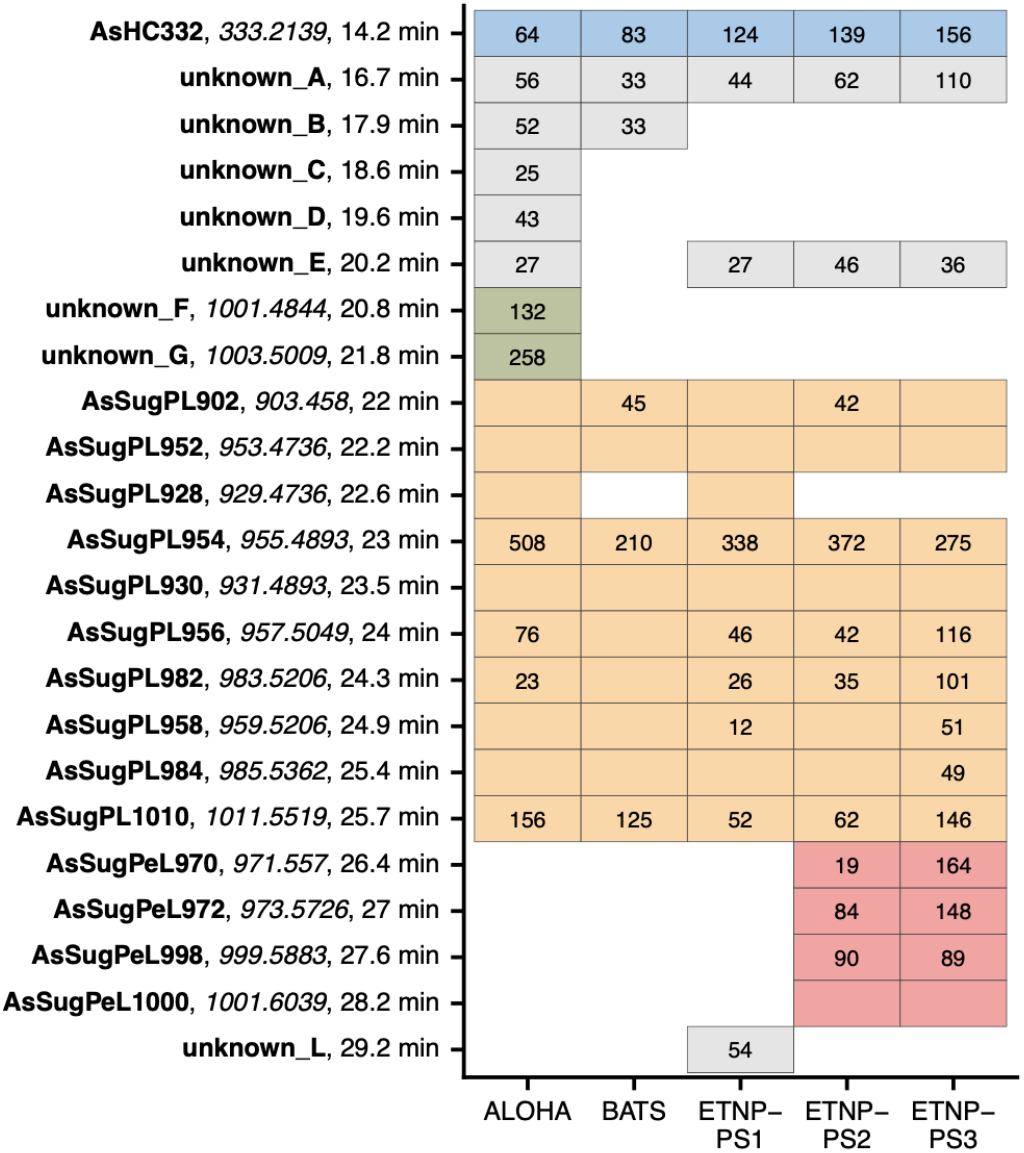
Arsenolipid quantification and characterization in each sample. Numbers within tiles are fmol L^−1^. Filled tiles are instances where the lipid was observed in the LC-HR-ESI-MS data, with values where we could quantify distinct peaks within the ^75^As LC-ICP-MS traces. Lipids are ordered by retention time and colored as in Figure 1.

We observed additional peaks in the ^75^As LC-ICP-MS chromatograms that we could not identify by searching for MS^1^ masses of known arsenolipids from our in-house database. In the oligotrophic ALOHA sample, we observed two large ^75^As LC-ICP-MS peaks at approximately 21 and 22 minutes and found two masses (1001.485 and 1003.500) with differing *m/z* and separation in retention time corresponding to one unsaturation at those times (labeled ‘unknown but *m/z* known’ in Figure 1 and Figure S1). Both masses had major MS^2^ fragments of 255.021, 237.010, and 195.000 *m/z* (Figure 3E and S1), which are likely associated with ions of C_7_H_16_O_5_As, C_7_H_14_AsO_4_, C_5_H_12_O_3_As, matching observed fragmentation spectra for other arsenosugar lipids (Raab et al. 2013; Glabonjat et al. 2018, 2019) but lacking fragments that would support a phosphate moiety. The high masses of these lipids (*m/z*’s > 1000) suggested two chains, but the retention times suggested that these lipids were more polar than the smallest observed AsSugPL.

In the ETNP-PS2 and ETNP-PS3 samples, we saw late eluting peaks in the LC-ICP-MS ^75^As chromatograms that did not correspond to any masses initially in our database (retention time > 26 minutes in Figure 1). After consideration, we have identified these as arsenosugar phospholipids with mixed acyl ether glycerols (AEG), which we refer to as arsenosugar phospho acyl ether lipids (AsSugPeL, Figure 2). We have four tiers of evidence to support these identifications. In the LC-HR-ESI-MS data, we saw near identical MS^2^ fragmentation between these lipids and the earlier eluting AsSugPL’s in the < 450 *m/z* range, strongly suggesting that these lipids all have the same headgroup (Figure 3 and S2). The masses that were isolated for fragmentation which resulted in these diagnostic fragments (971.55, 973.57, 999.59, and 1001.60) did not match any predicted *m/z*’s of AsSugPLs with either even or odd chain length but matched well to the calculated masses of AsSugPeLs (< 2 ppm for all except AsSugPeL972 which was < 6 ppm, Table S5). We also saw less abundant fragments in the MS^2^s of each of these lipids that correspond to the loss of the arsenosugar phospho headgroup of the lipid (neutral loss of 408.016) and to the loss of the fatty acid chain (*m/z* = 719.347, Figures 2 and S2). Finally, the later retention time relative to the AsSugPLs of similar masses suggests a less polar lipid class, consistent with an AEG. The even numbered nature on the ether-bound chain of the proposed structures suggest that the ether bound chains are not branched, though this could not be confirmed by our analyses.

## Discussion

### Bulk particulate arsenic measurements

Dissolved arsenic in open ocean surface waters is typically 10–18 nM (Cutter et al. 2001; Cutter and Cutter 2006). Our measurements show that particulate arsenic is nearly 1000 times less abundant than dissolved arsenic. This high seawater background obscures our view of transformations of arsenic occurring within or on particles and has hindered measurements of suspended particulate arsenic in aquatic systems for decades (Andreae 1978; Glabonjat et al. 2020). This background is important to take into consideration when loading particulate material onto a filter for analysis — we filtered 193–747 L of surface seawater onto 142 mm filter (Table S2). To account for sorbed dissolved arsenic we analyzed blanks which were never more than 10% of the signal observed in our samples. This work provides an important baseline for future particulate arsenic measurements, both in establishing a general range of values of open ocean particulate arsenic and providing methodology to obtain these data.

### Arsenolipids as part of the bulk particulate arsenic pool

Within the bulk particulate arsenic pool, we found that 7–20% was present in a lipid-soluble fraction (Table 1). This arsenolipid pool may act as an important conduit for arsenic transfer into the deeper ocean via sinking particles to maintain the nutrient-like profiles observed for this element (Cutter and Cutter 2006). Furthermore, arsenolipids have been hypothesized to be the precursor for arsenobetaine — an arsenic-containing metabolite that accumulates in mesopelagic particles and higher trophic levels (Duncan et al. 2015; Heal et al. 2020).

Lipid-bound arsenic was first observed in seaweed in the 1950s (Vinogradov 1953) and has since been seen within microalgae, macroalgae, and many species of fish (i.e. Lund 1973; Amayo et al. 2013; Raab et al. 2013; Taleshi et al. 2014; Xue et al. 2014; Glabonjat et al. 2018). Algal and cyanobacterial monocultures vary widely in the percent of total cellular arsenic present as arsenolipids in large part because of accumulated or sorbed inorganic arsenic (Duncan et al. 2013; Xue et al. 2014; Glabonjat et al. 2018, 2020). Our observations of 7–20% of total particulate arsenic as arsenolipids is on the higher end of these observations. Lund (1973) found that algae produced more arsenolipids under low arsenic conditions, which has also been observed in the cyanobacterium *Synechocystis* (Xue et al. 2014). These authors hypothesized that when arsenic exposure is low, arsenolipids serve as a biologically useful molecule (presumably a membrane lipid) and simultaneously act as a sink for arsenic, keeping the element’s cytosolic concentration low. We observed a higher percentage of particulate arsenic as arsenolipids in the ETNP samples (each about 20% of total particulate arsenic) than either oligotrophic samples (ALOHA and BATS). The ETNP samples are influenced by upwelling and likely have a higher dissolved phosphorous to arsenic ratio (P:As; and therefore lower arsenic exposure), hinting that arsenolipids may accumulate to higher levels when low arsenic exposure does not necessitate an immediate need to efflux toxic levels of arsenic.

### Composition of arsenolipids

In our samples, AsSugPLs were the most abundant type of arsenolipids. The diacyl varieties of these lipids have been observed in one species of cyanobacteria and several species of brown macroalgae (García-Salgado et al. 2012; Raab et al. 2013; Xue et al. 2014), thus we hypothesize that cyanobacteria or microbes related to brown macroalgae (e.g. diatoms) synthesize these lipids in the open ocean. No other marine observations of arsenolipids have been made, but recent work has characterized arsenolipids in sediments and suspended matter in relatively arsenic-rich lakes like Mono Lake in California (Glabonjat et al. 2020) and the Great Salt Lake in Utah (Glabonjat et al. 2019). At both locations, arsenolipid composition was dominated by a rich array of isoprenoidal arsenolipids related to phytol, hypothesized to be produced by green algae including *Picocystis* which dominate the Mono Lake phytoplankton community (Glabonjat et al. 2020). Additional green algae species produce arsenolipids of similar structure (Glabonjat et al. 2018; Řezanka et al. 2019). In contrast to these lake systems, we did not detect any of these isoprenoidal arsenolipids in these marine samples. Although chemotaxonomy of arsenolipids may be premature at this point, our observations do not support green algae as a major source for arsenolipids in the open ocean.

### Mixed acyl ether glycerol arsenolipids

Our observations of AEG arsenolipids is especially intriguing. To our knowledge, these are the first observations of ether-bound arsenolipids. Other AEGs have been seen in natural marine systems either as intact lipids (Schubotz et al. 2018) or after hydrolysis as mono alkyl glycerol ethers (MAGEs) (Hernandez-Sanchez et al. 2014). Although MAGEs have been proposed as a biomarker for anaerobic sulfate reducing bacteria (Orphan et al. 2001), MAGEs occur in other oxic open ocean sites (Hernandez-Sanchez et al. 2014), and other intact polar AEGs have been seen in surface waters near our study sites in the ETNP (Schubotz et al. 2018). Therefore, we think it is unlikely that our observed AEGs originate from sulfate-reducing bacteria in our oxic surface samples. Overall, while AEG lipids are unusual in surface waters, they are not completely unprecedented.

Most of the literature suggest eukaryotic algae as the main source of arsenolipids in natural systems (Glabonjat et al. 2019, 2020). Our provocative findings of AsSugPeL lipids suggest at least some of the lipids may be of bacterial origin. Ether-bound lipids in the surface ocean are hypothesized to originate from aerobic bacteria (Hernandez-Sanchez et al. 2014) and previous work in this area has hypothesized cyanobacteria as a possible source of AEGs after observing AEGs with a sulfoquinovosyl headgroup commonly associated with cyanobacteria (Schubotz et al. 2018). Especially at the ETNP-PS2 (offshore) and PS3 (coastal) stations where 20–30% of the identified arsenolipids were AEG lipids (Table 1, Figure 1), we hypothesize that bacteria — either cyanobacteria or aerobic heterotrophic bacteria — process arsenic in ways beyond efflux-based detoxification and biosynthesize arsenolipids.

## Conclusions

We analyzed marine suspended particles from five ocean locations to establish a baseline for particulate arsenic and its accumulation into lipids. This work supports the hypothesis that arsenolipid biosynthesis is a widespread phenomenon in marine systems. The characterization of the arsenolipid pool suggests bacteria as a possible overlooked source of these fascinating compounds in natural systems.

## Supporting information

Supplemental Tables, combined

## Acknowledgements

We thank A. Myers, L. Wood, L. Turner, T. Ugari, and J. Park for assistance with arsenolipid analysis; X. Zhang, S. Hayes, and members of the Zhang group for guidance, sampling equipment, and cruise prep assistance; S. Oleynik and K. Luxem for EA assistance; and L. Sall (funded by Princeton Environmental Institute summer internship) for assistance with EA sample prep. ETNP sampling was made possible by Chief Scientist B. Ward and the scientists and crew aboard the *R/V Sally Ride* cruise SR1805, funded by NSF Grant #1657663. BATS samples were provided by Y. Ryu and V. Luu aboard *R/V Atlantic Explorer* cruise AE1912, made possible by D. Sigman, funded by NSF grant #1536368. ALOHA sampling was made possible by M. Church, A. White, and the scientists and crew of the *R/V Kilo Moana* cruise KM1910 funded by NSF grant #1911831. This work was funded by the Simons Foundation (732763 to AEM, 598819 to KRH, 426570 to RMB and AEI, 385428 to AEI) and Princeton University Department of Geosciences Harry Hess Postdoctoral Fellowship (to AEM).

## Author Contributions

*KRH developed hypotheses, designed the study, adapted methodology, performed the laboratory and data analyses, and wrote the manuscript. KRH and AEM conducted field work. KRH and RMB performed data acquisition. KRH and AEI interpreted spectra. AEM provided ancillary data and additional samples. AEI and RMB provided access, materials, and instrumentation for analyses. KRH, AEM, AEI, and RMB edited and revised the manuscript*.

## Notes

### Competing Interest Statement

The authors have declared no competing interest.

