## Supplemental methods and figures for "Diverse arsenic-containing lipids in the surface ocean"

*Sample collection*. To sample suspended particles, we connected in-line filters in series followed by a flow meter and allowed the pressure from the clean underway intake to push water through the filters or used an air-powered PTFE diaphragm pump (Husky 307) connected to a hose submerged to a depth of 1–3 m (collection method noted in Table S1). We used an acid-washed 47 mm nylon prefilter (53 $\mu$m, Nitex brand) and a pre-combusted 142 mm GF-75 (nominal pore size 0.3 $\mu$m, Advantec) or 142 mm GF/F (nominal pore size 0.7 $\mu$m, Whatman), as noted in Table S1. For a subset of samples, we connected a third filter with an additional 142 mm GF-75, which we used as a wet blank – this represents the dissolved arsenic that sorbs onto the filter from seawater that is not associated with particles. This wet blank never had more than 10% of the arsenic signal from the samples. After a few hours, we recorded the volume filtered (reported in Table S1) and pumped the filters dry. All samples were folded in half (loaded side in) into new combusted aluminum foil and stored at –20 ºC until extraction.

*Cleaning procedures*. New and combusted (500 ºC for 5 hr) glassware (Pasteur pipettes and autosampler vials) and new plastics were not found to be a source of arsenic contamination and were used directly without additional acid cleaning. Glassware that was reused between samples was rinsed five times in MQ water (< 18.2 Ω, EMD Millipore), soaked for 24 hr in 10% HCl (trace metal grade), rinsed five times in MQ, dried in a particle free hood (Class 100 HEPA-filtered hood), and finally combusted at 500 ºC for 5 hr. Teflon vessels (Jensen Inert Products, Nalgene, or Savillex, depending on size) were carefully cleaned in order to minimize inter-sample contamination. Directly after reconstitution, Teflon tubes and caps were rinsed five times with MQ and heated (closed) in 50% $HNO_3$ (trace metal grade) for 2–3 hours (110 ºC). After the $HNO_3$ step, they were rinsed five times in MQ, soaked (24 hr) in 10% HCl (trace metal grade), rinsed five times in MQ, and dried in a particle free hood.

*Digestion for total particulate arsenic*. For quantification of total particulate arsenic, we took quantitative punches from each sample and digested the punches using only $HNO_3$ (Planquette and Sherrell 2012). We chose this digestion procedure because arsenic is known to combine with HCl and HF to form $AsCl_3$ or $AsF_3$ (Bajo 1978; Potolokov et al. 2006), which are volatile species that are lost during dry-down steps before analysis by mass spectrometry. Punches were placed in Teflon 15 mL flat bottomed jars (Savillex) and digested in 1 mL 50% $HNO_3$ at 110 ºC. After 4 hrs, samples were uncapped and the heat plate was turned up to 120 ºC to evaporate liquid. Once the samples were dried (approximately 30 min), they were removed from the heat plate and allowed to cool (capped). Once cooled, we added 1 mL of reconstitution solvent (2% $HNO_3$, 2.5% methanol, 1 ppb Rh in MQ water), heated the capped samples for 1 hr at 60 ºC to aid in solubilization, and syringe filtered samples into 2 mL glass autosampler vials (VWR). For each batch of digestions, we also digested a process blank (no sample) and at least one reference material (Kelp *Thallus laminariae* SRM3232 from NIST or Cuttlefish SQID-1 from Canadian reference materials) to monitor for digestion efficacy and reagent contamination. The 2.5% methanol in the reconstitution solvent masked any changes in $^{75}As$ signal associated with carbon loading, a known phenomenon (Larsen and Stürup 1994).

*Extraction for arsenolipids*. To extract for arsenolipids, we used approximately half of each sample on 142 mm GF/F or GF-75 filter, some with punches removed. See Table S1 for effective extraction volumes. We separated the filter into two 50 mL Teflon tubes with screw top Teflon lids (Nalgene) and submerged each filter in 10 mL of 2:1 dichloromethane:methanol per tube – an extraction technique used by others investigating arsenolipids (Glabonjat et al. 2014). We then sonicated the tubes with samples and solvent for 10 min, followed by shaking on a shaker table for 10 min at 120 rpm. We repeated this sonication and shaking three times for a total extraction time of 60 min, with manual end-over-end mixing between each sonication and shaking step. After the physical disruption, we carefully decanted the solution into a 50 mL combusted glass centrifuge tube. The glass fiber filters used in collection of marine particles are particularly porous and do not allow for a complete transfer of the liquid from the Teflon tubes, so we added 10 mL more of the 2:1 dichloromethane:methanol and repeated the extraction process twice more. Overall, this extraction yielded about 30 mL of solution containing mostly lipid-soluble compounds. To clean up the lipid-soluble compounds of water-soluble arsenic species, we added 10 mL of water (MQ), centrifuged the mixture at 1200 rpm for 10 min, and removed the top layer (containing water and methanol) into a clean 50 mL combusted glass centrifuge tube. The remaining solution of dichloromethane was dried down under clean $N_2$ and reconstituted in 1.5 mL of 9:1 methanol:toluene.

To clean up the crude lipid fraction, we performed a silica gel clean up as previously described (Glabonjat et al. 2014). We prepared a glass Pasteur pipette (150 × 5 mm) with glass wool and 5 cm of silica gel 60. We conditioned the column with 10 mL of 1:1 methanol:acetone with 1% formic acid before loading 1 mL of our crude extract. We washed the column with 5 mL of 1:1 methanol:acetone with 1% formic acid and 5 mL methanol and then eluted the samples with 10 mL of methanol with 1% ammonia. This eluent was dried down under clean $N_2$ and reconstituted in 1 mL of 9:1 methanol:toluene.

*Details on liquid chromatography*. Chromatographic separation for both LC-ICP-MS and LC-HR-ESI-MS was performed on a Dionex Ultimate 3000 bio-inert LC, using a C18 column (ZORBAX SB-C18 by Agilent, 0.5x150 mm, 5 $\mu$m particle size, 80 Å pore size) maintained at 55 ºC with a flow rate of 40 $\mu$L min$^{-1}$. The low flow rate eliminated the need for post-column desolvation or dilution with water. We employed a gradient elution with 1% formic acid in water (solvent A) and 1% formic acid in isopropyl alcohol (IPA, solvent B). The gradient started at 30% B held for 5 min, ramped from 30% B to 90% B over 25 min, held at 90% B for 15 min, ramped back to 30% B over 3 min before re-equilibrating at 30% B for 7 min before the next injection. Crude extracts and silica-cleaned extracts were injected at 25 $\mu$L for all analyses; extracts were injected once in LC-ICP-MS configuration (for quantification), and twice in LC-HR-ESI-MS configuration (for characterization).

*Details on LC-ICP-MS analysis*. The ICP was equipped with platinum sampler and skimmer cones, a 1 mm quartz injector, an organics torch, a cyclonic spray chamber cooled to 0 ºC, and a perfluoroalkoxy microflow nebulizer equipped with exchangeable sample uptake capillary (PFA-ST, Elemental Scientific). We used $O_2$ as an additional gas in the spray chamber at 8% to aid in solvent combustion to prevent carbon deposits on the cones. We tuned the nebulizer gas flow and collision gas flow daily using an uptake capillary designed for flows at 50 $\mu$L min$^{-1}$ with a mixture of cobalt (as cyanocobalamin) and arsenic (as arsenobetaine) each at 250 nM at the starting chromatographic gradient conditions (70:30 water:IPA with 1% formic acid). This resulted in a nebulizer flow (Ar) of 0.74 mL min$^{-1}$ with a collision gas flow (He) of 2.0 mL min$^{-1}$

which eliminated $^{35}Cl^{40}Ar$ interference. We monitored the following masses at the respective dwell times: $^{75}As$ (0.25 s), $^{59}Co$ (0.03 s), $^{77}Se$ (0.03 s), $^{78}Se$ (0.01 s), and $^{31}P$ (0.01 s).

*Details of arsenolipid quantification*. There is no commercial standard for organic arsenic compounds that can be retained on a C18 column, so in order to calculate concentrations of the arsenolipids, we used a technique similar to previous work (Boiteau et al. 2016). This approach uses a proxy element and establishes the relative instrumental response of the proxy element to the analyte element by direct injection under chromatographic conditions. We ran a standard curve of cyanocobalamin at our chromatographic conditions to establish an instrument response to $^{59}Co$. We then intercalibrated signal sensitivities of $^{59}Co$ and $^{75}As$ by direct injection of a mixture of cobalt (as cyanocobalamin) and arsenic (as arsenobetaine) each at 250 nM at the starting chromatographic gradient conditions (30% solvent B, or 70:30 water:IPA with 1% formic acid). Since the $^{75}As$ is affected by carbon loading, we also injected arsenobetaine at 50%, 70%, and 90% solvent B to establish a correction factor to apply to the $^{75}As$ signal – overall we saw a 19% decrease in $^{75}As$ signal over the full LC gradient range (30% and 90% B), and we corrected for this decrease in signal when calculating concentrations over the course of the gradient.

*Details on LC-HR-ESI-MS data collection*. The LC-HR-ESI-MS contained a heated electrospray ionization source with a capillary temperature of 275 ºC , sheath gas of 8 (arbitrary units), auxiliary gas equal to 1 (arbitrary units), with no sweep gas. Spray voltage was set at 3.5 kV, auxiliary gas heater temperature of 130 ºC, and S-lens RF level of 65.0. To obtain higher quality $MS^1$ and $MS^2$ data, we injected each sample twice, once in a low mass configuration and once in a high mass configuration. For both LC-HR-ESI-MS configurations we used a data-dependent acquisition (DDA) with 10 s dynamic exclusion, with $MS^1$ scans collected at 60,000 resolution in positive mode from either 200–560 (low mass injections) or 650–1090 *m/z* (high mass injections). $MS^2$ spectra were collected using an isolation window of 0.4 *m/z* at 60,000 resolution from 70–560 *m/z* (low mass injections) or 70–1090 *m/z* (high mass injections). We employed two inclusion lists (one for low mass and one for high mass injections) of possible masses of arsenolipids based on literature searches and *in-silico* database construction (see below) in order to obtain $MS^2$ s of relevant $MS^1$s. $MS^2$ scans were collected at ramped 20, 35, and 50 normalized collision energies.

*Details on LC-HR-ESI-MS data analysis and arsenolipid identification*. To aid in our search for arsenolipids in our LC-HR-ESI-MS data, we constructed a database of possible arsenolipid structures based on previous studies. For the fatty acid and glycolipids we included theoretical masses for lipids with a wide range of chain lengths and saturations. We calculated predicted $MS^1$ spectra (based on M+H ionization, which is generally the observed ion) and major peaks in the $MS^2$ spectra. See Table S2 for description of the *in-silico* arsenolipid database including the literature used as a basis for $MS^1$ and $MS^2$ predictions, and Tables S3 and S4 for the full databases.

Using the $MS^1$ database described above (Table S3), we searched for corresponding signals in the LC-HR-ESI-MS data that corresponded to retention times where we observed a peak in the LC-ICP-MS $^{75}As$ signal. We searched for the monoisotopic mass of each lipid and also the $^{13}C$ isotope of each lipid (scaled to predicted $^{13}C$ /$^{12}C$ ratio), which gave us more confidence in the exact mass match. Next, we compared the $MS^2$ fragmentation to the expected (from literature) fragmentation to assign identifications to possible arsenolipids in the ICP-MS $^{75}As$ signal. Our assignments follow the expected retention times from literature well, with the earliest peaks corresponding to arseno-hydrocarbons, and later peaks corresponding to intact polar arsenosugar phospholipids.

To go beyond the results from the $MS^1$ search, we also searched the $MS^2$ scans for masses that were associated with arsenolipids and contained arsenic. Arsenic's negative mass defect and odd number allowed us to search for several masses in the $MS^2$ spectra that unequivocally contain arsenic. This yielded several additional assignments of the peaks in the LC-ICP-MS $^{75}As$ signal to a *m/z,* though not always to a complete identification.

**Supplemental Tables**

Table S1: Expanded sample descriptions including date collected, collection method, filter set-up, volume filtered (onto 142 mm filter), effective volume digested for bulk arsenic quantification, effective volume extracted for arsenolipid characterization, and the names of the four LC-HR-ESI-MS files that correspond to each sample. Those data files will be available on Dryad upon publication.

Table S2:
Summary of lipid structures included in search database and associated literature. For the more rarely seen AsIsop, AsPE, and TMAsFOH lipids we used only the exact formulae seen before in literature values rather than a complete set of iterations of possible lipids. Reference 1: Amayo et al. (2011); 2: Amayo et al. (2013); 3: Arroyo-Abad et al. (2016); 4: Lischka et al. (2013); 5: Pétursdóttir et al. (2018); 6: Raab et al. (2013); 7: Rumpler et al. (2008); 8: Sele et al. (2014); 9: Viczek et al. (2016); 10: García-Salgado et al. (2012); 11: Glabonjat et al. (2018); 12: Glabonjat et al. (2020); 13: Pétursdóttir et al. (2015); 14: Pétursdóttir et al. (2018); 15: Taleshi et al. (2008); 16: Taleshi et al. (2010); 17: Taleshi et al. (2014); 18: Xue et al. (2014)

Table S3:
Database for searching for arsenolipids in $MS^1$ LC-HR-ESI-MS data, as summarized in Table S2.

Table S4:
Database for searching for arsenolipids in $MS^2$ LC-HR-ESI-MS data, as summarized in Table S2.

Table S5:
Expected and observed *m/z* for AsSugPeL970, 972, 998, and 1000 in ETNP-P2, ETNP-P3, and ALOHA.

**Supplemental Figures:**

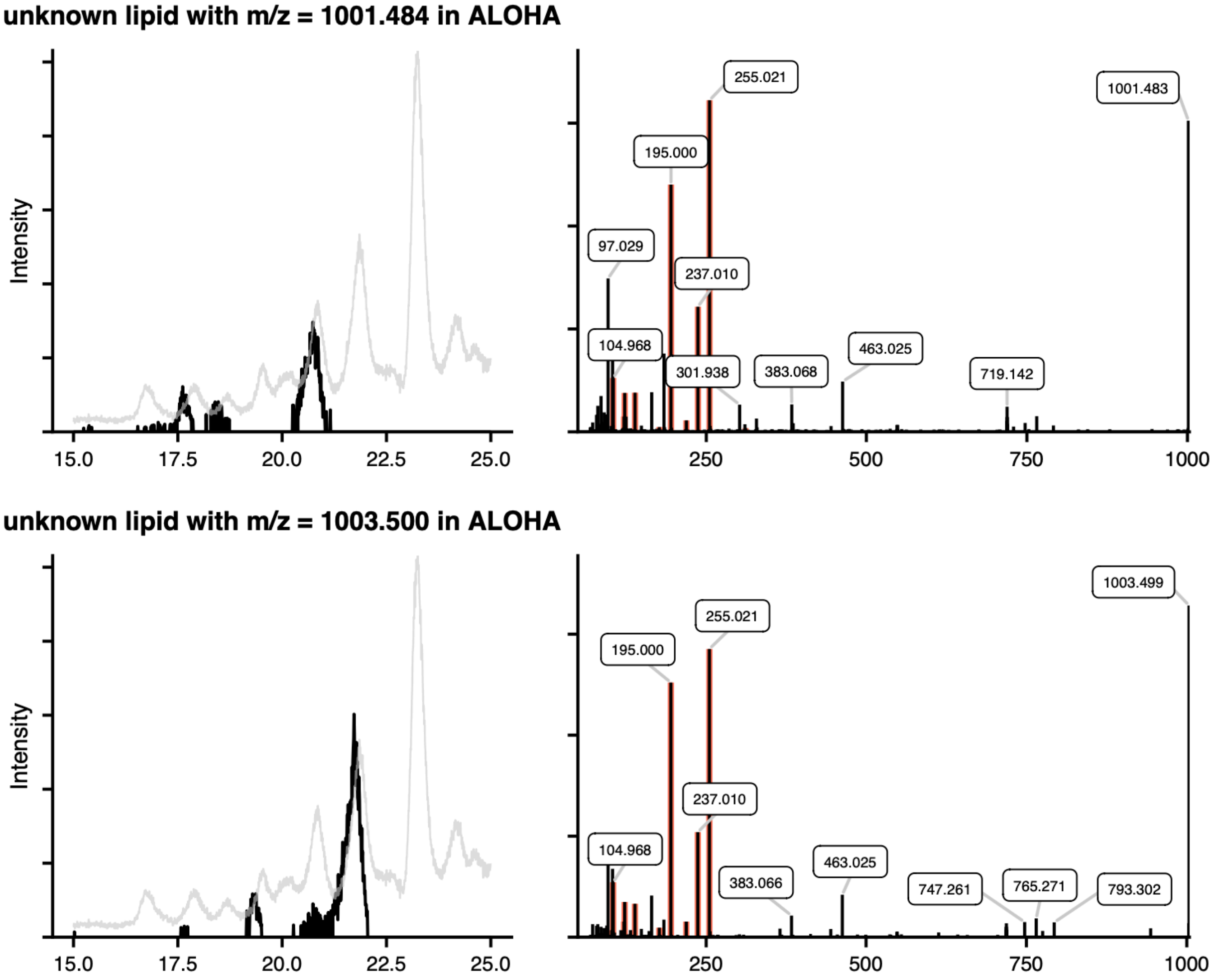


Figure S1: Unknown arsenolipids found in ALOHA sample. Left panels show extracted ion chromatograms (within 0.01 *m/z* of masses noted in figure) from LC-HR-ESI-MS (in black) overlaid on the LC-ICP-MS $^{75}As$ signal (in grey). Right panels show $MS^2$ spectra, with diagnostic arsenic masses highlighted and major masses labeled.

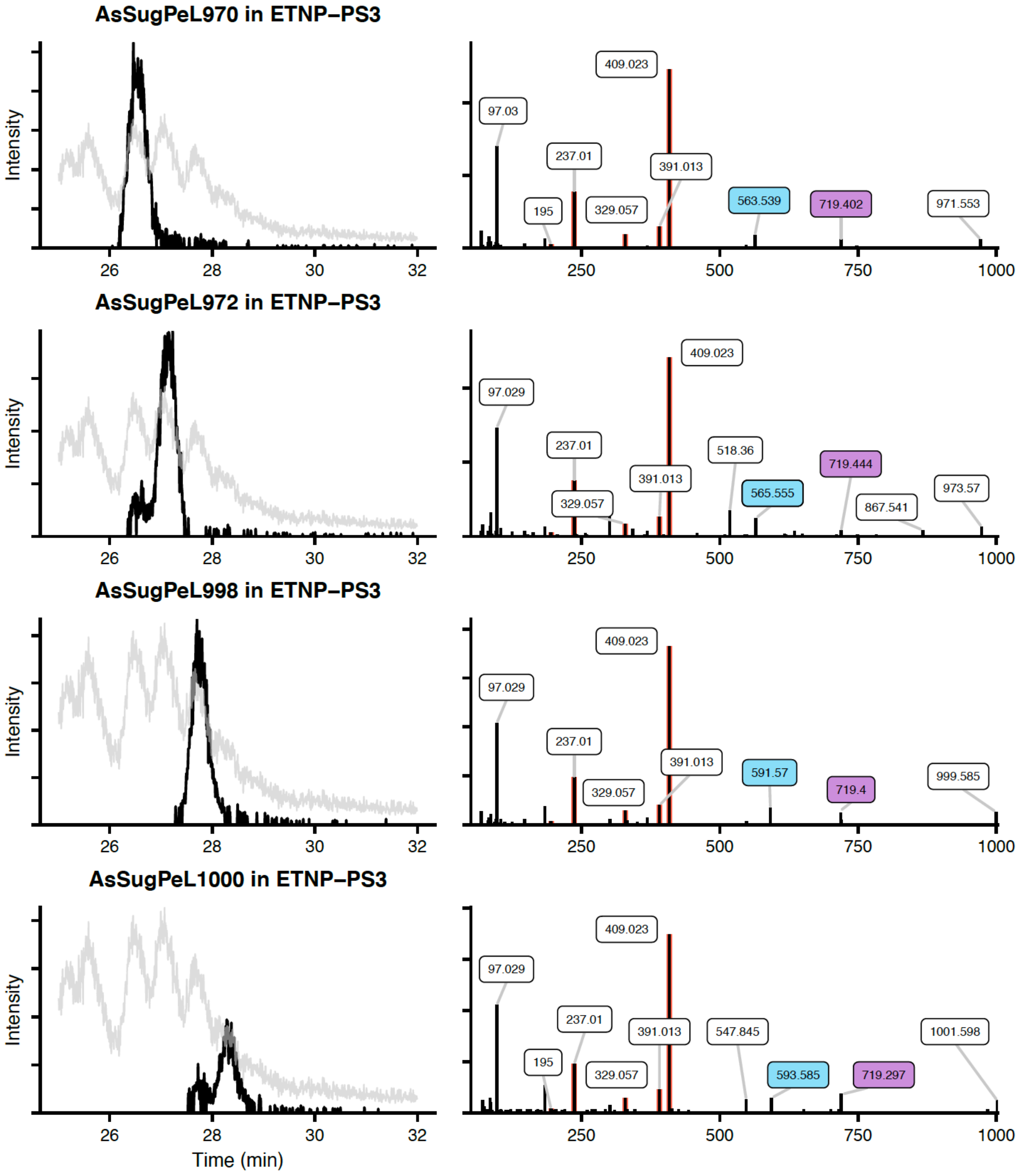


Figure S2: AsSugPeLs found in ETNP-PS3 sample. Left panels show extracted ion chromatograms for identified lipids from LC-HR-ESI-MS data (in black, at tolerance of $m/z$ = 0.01) overlaid on the LC-ICP-MS $^{75}$As signal (in grey). Right panels show $MS^2$ spectra, with diagnostic arsenic-containing masses highlighted in orange. Masses highlighted in blue correspond to neutral loss of the AsSugPL headgroup, masses in purple correspond to loss of the fatty acid.
